# Signal Transduction in Human Cell Lysate via Dynamic RNA Nanotechnology

**DOI:** 10.1101/439273

**Authors:** Lisa M. Hochrein, Tianjia J. Ge, Maayan Schwarzkopf, Niles A. Pierce

## Abstract

Dynamic RNA nanotechnology with small conditional RNAs (scRNAs) offers a promising conceptual approach to introducing synthetic regulatory links into endogenous biological circuits. Here, we use human cell lysate containing functional Dicer and RNases as a testbed for engineering scRNAs for conditional RNA interference (RNAi). scRNAs perform signal transduction via conditional shape change: detection of a subsequence of mRNA input X triggers formation of a Dicer substrate that is processed to yield siRNA output anti-Y targeting independent mRNA Y for destruction. Automated sequence design is performed using the reaction pathway designer within NUPACK to encode this conditional hybridization cascade into the scRNA sequence subject to the sequence constraints imposed by X and Y. Because it is difficult for secondary structure models to predict which subsequences of mRNA input X will be accessible for detection, here we develop the RNAhyb method to experimentally determine accessible windows within the mRNA that are provided to the designer as sequence constraints. We demonstrate the programmability of scRNA regulators by engineering scRNAs for transducing in both directions between two full-length mRNAs X and Y, corresponding to either the forward molecular logic “if X then not Y” (X ┤ Y) or the reverse molecular logic “if Y then not X” (Y ┤ X). In human cell lysate, we observe a strong OFF/ON conditional response with low crosstalk, corresponding to a ≈20-fold increase in production of the siRNA output in response to the cognate vs non-cognate full-length mRNA input. Because diverse biological pathways interact with RNA, scRNAs that transduce between detection of endogenous RNA inputs and production of biologically active RNA outputs hold great promise as a synthetic regulatory paradigm.

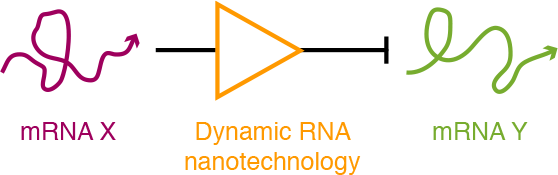

---

Over the last 15 years, researchers in the emerging discipline of dynamic DNA nanotechnology have developed striking capabilities for engineering pathway-controlled hybridization cascades in which small conditional DNAs (scDNAs) execute dynamic functions by autonomously performing interactions and conformation changes in a prescribed order.^1, 2^ These mechanisms are powered by the enthalpy of base pairing and the entropy of mixing, exploiting diverse design elements to effect assembly, disassembly, and pathway control (Figure 1a). While considerable effort has been invested in exploring dynamic DNA nanotechnology in a test tube, comparatively little attention has been paid to dynamic RNA nanotechnology, which offers profound opportunities for introducing synthetic regulatory links into living cells and organisms. We envision small conditional RNAs (scRNAs) that interact and change conformation to transduce between detection of an endogenous programmable input, and production of a biologically active programmable output recognized by an endogenous biological pathway (Figure 1b). In this scenario, the input controls the scope of regulation and the output controls the target of regulation, with the scRNA leveraging programmability and conditionality to create a logical link between the two.

As a motivating example, consider conventional RNA interference (RNAi), which offers the benefit of programmability, but lacks conditionality. RNAi mediated by small interfering RNAs (siRNAs) enables knockdown of a gene of choice,^3, 4^ executing the molecular logic: silence gene Y. Because siRNAs are constitutively active, it is difficult to confine silencing to a subset of the cells under study (e.g., Figure 1c top). To enable cell-selective silencing, we envision scRNAs that mediate “conditional RNAi” corresponding to the molecular logic: if gene X is transcribed, silencing independent gene Y. Upon detection of mRNA input X, the scRNAs perform shape and sequence transduction to form a Dicer substrate that is processed by Dicer to yield siRNA output anti-Y targeting an independent mRNA Y for destruction. In this scenario, Y would be silenced only in tissues where X was expressed (e.g., Figure 1c bottom).

Several groups have made contributions^5–9^ toward the still outstanding goal of engineering scRNAs that perform shape and sequence transduction to implement conditional RNAi (X ┤ Y). In this endeavor, we previously developed five scRNA mechanisms for conditional Dicer substrate formation, each exploiting a different combination of design elements from Figure 1a.^8^ In the presence of full-length mRNA input X, these mechanisms produce an OFF/ON conditional response yielding an order of magnitude increase in production of a Dicer substrate targeting independent mRNA Y. By appropriately dimensioning and/or chemically modifying the scRNAs, only the product of signal transduction, and not the reactants or intermediates, was efficiently processed by Dicer, yielding siRNA outputs anti-Y. In broad terms, this work demonstrated that design elements previously developed and explored in the context of dynamic DNA nanotechnology could be adapted for dynamic RNA nanotechnology, providing a molecular vocabulary for pursuing the regulatory goals of Figure 1b.

The simplest and most promising of these mechanisms employed a duplex scRNA A·B, which upon detecting mRNA input X, produced an shRNA Dicer substrate B that is processed by Dicer to yield siRNA output anti-Y (Figure 1d).^8^ With this scRNA mechanism, X partially displaces A from B via toehold-mediated 3-way branch migration (Step 1a), exposing a previously sequestered internal toehold, ‘c’, within B, mediating a further 3-way branch migration (Step 1b) that promotes spontaneous dissociation of B from X · A to yield shRNA Dicer substrate B. This mechanism has the desirable property that the scRNA A · B is stable rather than metastable (i.e., if scRNAs are allowed to equilibrate in the absence of X, they will predominantly remain in the A · B reactant state rather than producing product shRNA B).^8^ Hence, there is a thermodynamic rather than a kinetic limit on the amount of spurious output that will be produced in the absence of input, providing the conceptual basis for engineering a clean and reliable OFF state. When we measured the OFF/ON conditional response for production of shRNA B in a test tube, the OFF state was undetectable (i.e., smaller than the gel quantification uncertainty), illustrating the benefits of using a stable scRNA. The OFF/ON conditional response with a full-length mRNA input X was >50-fold. On the other hand, using a short RNA input X_s_ corresponding to only the portion of the mRNA that binds the scRNA, the OFF/ON ratio was >200-fold, suggesting that the secondary structure of the full-length mRNA inhibited interactions with the scRNA to some degree.^8^

**Figure 1.**
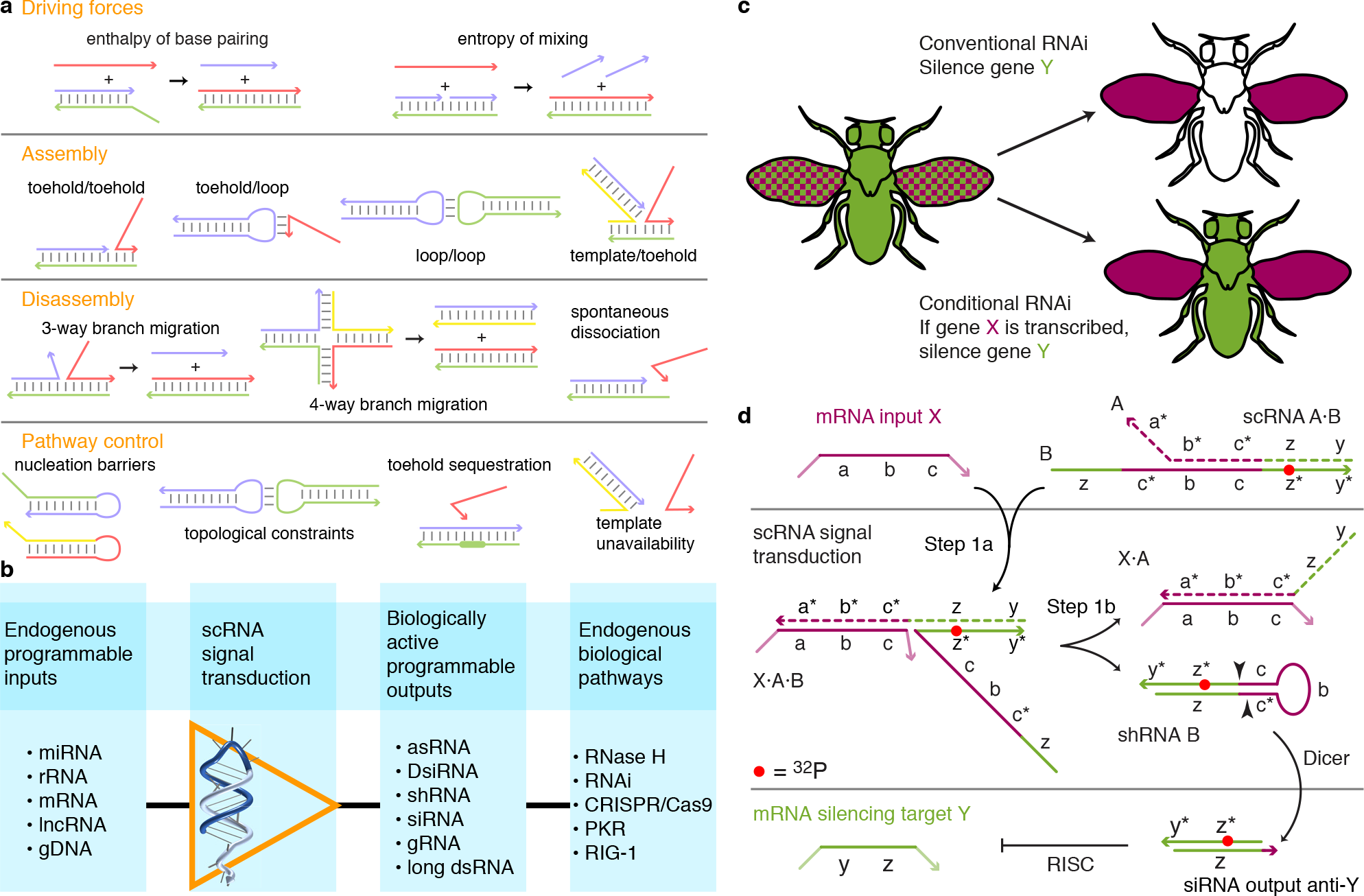
Dynamic RNA nanotechnology for programmable conditional regulation with small conditional RNAs (scRNAs). (a) Design elements for dynamic RNA nanotechnology. (b) Opportunities for programmable conditional regulation via scRNA signal transduction. (c) Molecular logic of conventional RNAi (“not Y”; ┤ Y) vs. conditional RNAi (“if X then not Y”; X ┤ Y). In this conceptual illustration, conventional RNAi silences Y in all tissues, while conditional RNAi silences Y only in tissues where and when X is expressed, exerting spatiotemporal control over regulation. (d) scRNA mechanism implementing molecular logic X ┤ Y. scRNA A·B detects mRNA input X (containing subsequence ‘a-b-c’), leading to production of shRNA Dicer substrate B that is processed by Dicer to produce siRNA output anti-Y (targeting mRNA silencing target Y containing subsequence ‘y-z’). scRNA A·B is stable in the absence of X. X partially displaces A from B via toehold-mediated 3-way branch migration, exposing a previously sequestered internal toehold, ‘c’, within B (Step 1a). Internal nucleation of duplex ‘c/c*’ within B mediates a further internal 3-way branch migration to form duplex ‘z/z*’, facilitating disassembly of B from X·A via spontaneous dissociation of short duplex ‘y/y*’ (Step 1b). Dicer processing of the resulting canonical shRNA Dicer substrate, B, yields siRNA output anti-Y (with guide strand ‘z*-y*’). Domain lengths: |a| = 16, | b| =14, | c| = 5, |y| = 2, |z|=19. RNA strands denoted by uppercase letters, sequence domains denoted by lowercase letters with complementarity indicated by ‘*’. ^32^P internal radiolabel denoted by dot on B strand. Chemical modifications (2′OMe-RNA): A (dashed backbone), B (5′ and 3′ nucleotides only).

Here, as a stepping stone toward validating scRNA signal transduction in living cells and organisms, we tested this scRNA mechanism in human cell lysate. Compared to our previous test tube studies in buffer using recombinant Dicer, lysate with functional endogenous Dicer^10^ more closely mimics the cellular environment, as it includes cellular proteins and nucleic acids that can inhibit scRNA function, and RNases that can degrade the scRNAs (we specifically avoided adding RNase inhibitors to the lysate in order to preserve the challenge of overcoming RNase-mediated degradation). Notably, working in lysate eliminates the need to deliver scRNAs across the cell membrane, removing a potential failure mode in order to focus attention on the performance of the signal transduction mechanism itself.

Our previous experiments were performed using scRNAs and full-length mRNA inputs at 0.5 *μ*M in buffer, assayed with native polyacrylamide gel electrophoresis and fluorescent staining. Because previous RNAi studies suggest the need for an siRNA concentration in the range of 0.1-10 nM within the cell for effective gene knockdown in vivo,^11, 12^ we set out to test scRNA A · B at 2.5 nM in our lysate studies. To provide the sensitivity needed for gel assays at this low concentration, as well as to discriminate between scRNA-derived and lysate-derived nucleic acids, we radio-labeled strand B with ^32^P (Section S1.2), enabling tracking of state changes between: 1) duplex scRNA A · B, 2) monomer shRNA Dicer substrate B, 3) Dicer-processed siRNA output anti-Y, and 4) RNase-mediated degradation of B. The full-length mRNA input X (or short RNA input X_s_) was introduced at 4× the scRNA concentration. With scRNAs at the low 2.5 nM concentration in buffer, we observed strong conditional production of shRNA B in response to short RNA input X_s_ but minimal response to the full-length mRNA input X (Figure S4a). Furthermore, in human cell lysate, the response to the short RNA input X_s_ was substantially weakened, and there was negligible response to the full-length mRNA input X (Figure S4b). These initial failures provided the embarkation point for the present work, in which we use human cell lysate as an engineering testbed to optimize scRNA performance for full-length mRNA detection at low concentration in the presence of cellular proteins, nucleic acids, and RNases.

These initial results suggested the need to re-dimension the scRNA sequence domains to optimize the energetics of the mechanism for detection of full-length mRNA input X at low concentration in lysate. Over several rounds of sequence design and experimental testing, we made the following adjustments to scRNA A·B: 1) increased toehold ‘a’ by 4 nt to enhance nucleation between A·B and X and better mediate subsequent partial opening of duplex A · B via toeholdmediated 3-way branch migration (Step 1a), 2) increased duplex ‘c/c*’ by 2 bp to enhance self-nucleation within B and better mediate subsequent further opening of duplex A·B via toehold-mediated 3-way branch migration (Step 1b), 3) addition of 2-nt duplex ‘y/y*’ to A·B to require spontaneous dissociation of B from A·B to reduce spurious production of shRNA B in the absence of input mRNA X.

For each design cycle with new scRNA domain dimensions, sequence design was performed using the reaction pathway designer within NUPACK;^13, 14^ sequence design was formulated as a multistate optimization problem using target test tubes to represent reactant, intermediate, and product states along the reaction pathway (Figure 2a).^14^ Each target test tube contains the depicted on-target complexes corresponding to the on-pathway products for a given step (each with the depicted target structure and a target concentration of 2.5 nM) as well as off-target complexes (all complexes of up to 4 strands, each with a target concentration of 0 nM; not depicted) corresponding to on-pathway reactants and offpathway crosstalk for a given step.

Sequence design is performed subject to complementarity constraints inherent to the reaction pathway (Figure 1d; domain ‘b’ complementary to ‘b*’, etc), as well as to biological sequence constraints imposed by the mRNA input X and the mRNA silencing target Y. For the current scRNA mechanism, every nucleotide in the design is constrained to be a subsequence of either X or Y (see constraint shading in Figure 2a), reflecting the simplicity of the mechanism, and placing severe demands on sequence design.

The sequence is optimized by reducing the ensemble defect, quantifying the average fraction of incorrectly paired nucleotides over the multi-tube ensemble.^14, 17, 18^ Optimization of the ensemble defect implements both a positive design paradigm, explicitly designing for on-pathway elementary steps, and a negative-design paradigm, explicitly designing against off-pathway crosstalk. The ensemble defect can be decomposed into two types of contribution: “structural defect” (fraction of nucleotides in the incorrect basepairing state within the ensemble of the on-target complex) and “concentration defect” (fraction of nucleotides in the incorrect base-pairing state because there is a deficiency in the concentration of the on-target complex).

Figure 2b displays the target test tubes for a completed sequence design of the redimensioned scRNA A · B. Concentration defects are negligible (each on-target complex forms with quantitative yield at the desired 2.5 nM target concentration). On the other hand, the structural defect is above 6% for scRNA A · B (see probability shading in Figure 2b indicating undesired base-pairs between toeholds ‘a*’ and ‘z’ with probability greater than zero) and for intermediate B_truncate_ (indicating desired base-pairs in duplex ‘c/c*’ with probability less than one). These defects reflect the real-world challenges of designing sequences that execute a dynamic reaction pathway, yet are fully constrained by the sequences of mRNAs X and Y.

**Figure 2.**
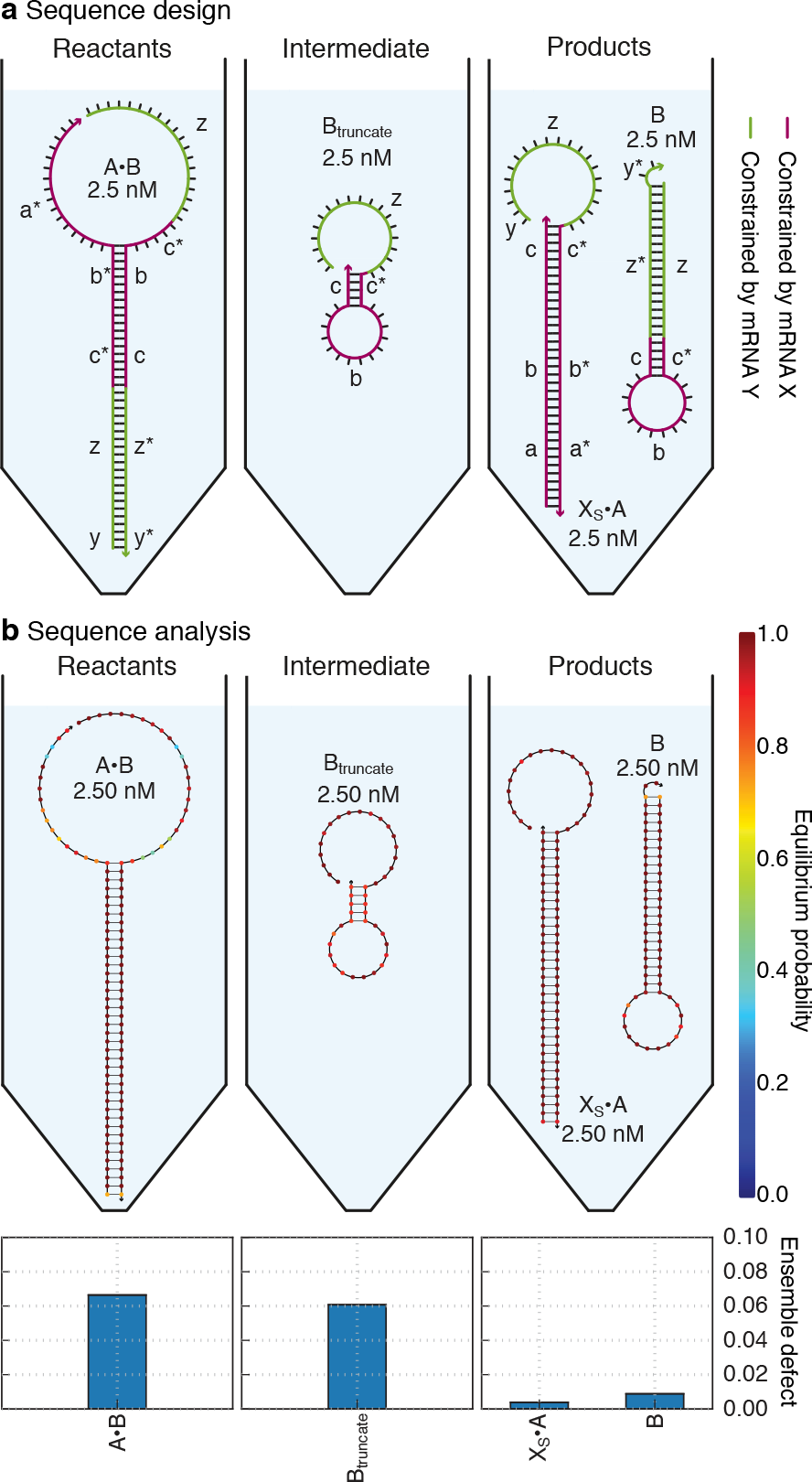
Computational scRNA sequence design using NUPACK. (a) Sequence design is formulated as a multistate optimization problem using target test tubes to represent reactant, intermediate, and product states along the reaction pathway (Figure 1d).^13, 14^ Each target test tube contains the depicted on-target complexes corresponding to the on-pathway products for a given step (each with the depicted target structure and a target concentration of 2.5 nM) as well as off-target complexes (all complexes of up to 4 strands, each with a target concentration of 0 nM; not depicted) corresponding to on-pathway reactants and off-pathway crosstalk for a given step. The Intermediate tube (Step 1a) contains a truncated version of strand B to facilitate design of the ‘c/c*’ self-nucleation duplex. Every nucleotide in the design is constrained by the sequence of either the mRNA input X or the mRNA silencing target Y (see domain shading). (b) Analysis of design quality over the design ensemble.^13, 15^ Tubes depict the predicted concentration and target structure for each on-target complex, with nucleotides shaded to indicate the probability of adopting the depicted base-pairing state at equilibrium. For this design, all on-targets are predicted to form with quantitative yield at the 2.5 nM target concentration (negligible concentration defects) but some nucleotides have unwanted base-pairing interactions (non-negligible structural defects for nucleotides not shaded dark red). Bar graphs depict the residual defect for each on-target complex in each tube (blue shading: structural defect component, green shading: concentration defect component [negligible for this sequence design]). RNA at 37 °C in 1 M Na+.^16^

**Figure 3.**
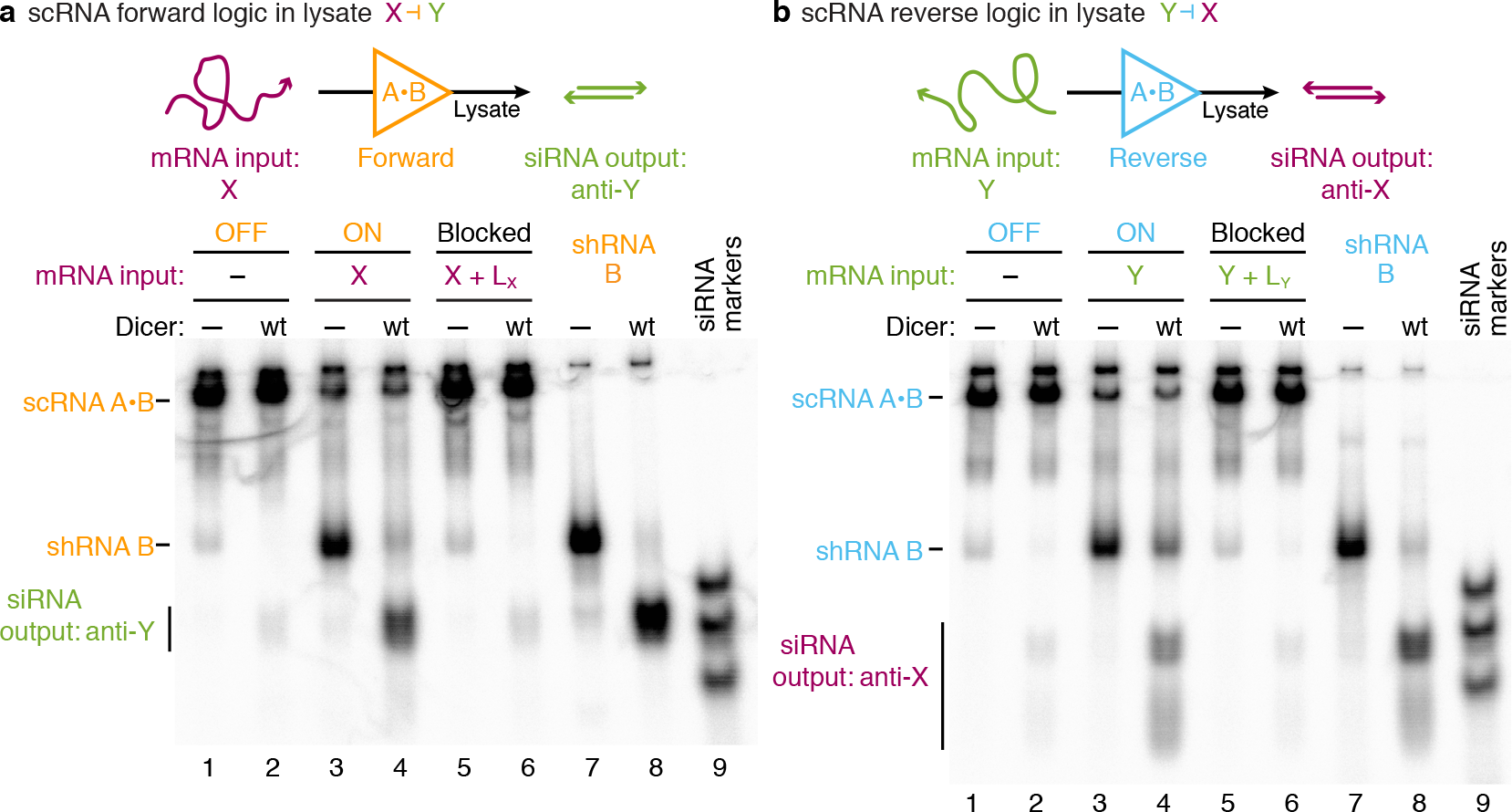
Conditional siRNA production via scRNA signal transduction in human cell lysate. (a) Forward molecular logic (X ┤ Y): if mRNA input X is detected, generate siRNA output anti-Y targeting mRNA Y for silencing. (b) Reverse molecular logic (Y ┤ X): if mRNA input Y is detected, generate siRNA output anti-X targeting mRNA X for silencing. (a,b) scRNA signal transduction in lysate (HEK 293T) that is either Dicer-depleted (—) or wildtype (wt; containing endogenous Dicer). Strand B internally radiolabeled with ^32^P (dot in Figure 1d) to enable native PAGE assay with scRNA at 2.5 nM; mRNA input spiked into lysate at ≈10 nM. OFF state: minimal production of siRNA output in the absence of mRNA input (lanes 1, 2) or in the blocked state where the mRNA input is pre-incubated with a blocker strand L that binds to the scRNA nucleation site on the mRNA (lanes 5, 6). ON state: strong production of siRNA output in the presence of mRNA input (lanes 3, 4). shRNA B provides a control illustrating Dicer processing to generate siRNA output (lanes 7, 8). siRNA size markers (lane 9).

Operating at 2.5 nM in human cell lysate, the redimensioned scRNA exhibits strong OFF/ON conditional siRNA production (Figure 3a). In the absence of mRNA input X, there is minimal production of siRNA output anti-Y (lane 2), while in the presence of X, there is strong production of anti-Y (lane 4). To verify that siRNA production is Dicer mediated, we depleted Dicer from the lysate (Dicer knockdown in cells using an siRNA pool followed by lysate immunodepletion with an anti-Dicer antibody). The Dicer-depleted lysates show minimal siRNA production (lanes 1 and 3) relative to wildtype lysates (lanes 2 and 4), producing shRNA B instead of siRNA anti-Y as the product of signal transduction. To verify that A ┤ B signal transduction is triggered by the intended subsequence of mRNA input X, we pre-incubated X with an 2′OMe-RNA blocker strand, LX, that binds to the nucleation site, ‘a’, on X (lanes 5 and 6), effectively blocking interaction with A·B and restoring the OFF state (cf. lanes 1 and 2).

The data of Figure 3a demonstrate conditional siRNA production for the forward molecular logic “if X then not Y” (X ┤ Y), where X is a full-length mRNA input and Y is an independent mRNA silencing target. With the intention of demonstrating the programmability of scRNA signal transduction, we repeated sequence design for the reverse molecular logic “if Y then not X” (Y ┤ X), using Y as the mRNA input and X as the mRNA silencing target. For multiple sequence designs, we ran into the difficulty that the designed scRNAs functioned properly in detecting the short RNA input Y_s_ (the targeted subsequence of Y), but were ineffective in detecting the full-length mRNA input Y (see two examples in Figures S5 and S6). This failure mode highlights the inherent difficulty in attempting to engineer scRNAs that detect full-length mRNAs. The RNA secondary structure model^16^ underlying NUPACK is not as accurate for predicting structural properties of (long) mRNAs as for (short) scRNAs, presumably due to a combination of pseudoknotting, long-range tertiary interactions, and (in lysate) protein/RNA interactions − none of which are accounted for in the physical model. In effect, the sequence design pipeline succeeds in engineering the portion of the problem that is present in the model, but it fails to overcome the challenges of mRNA structure that are omitted from the model.

After repeatedly encountering this problem while attempting to engineer the reverse logic Y ┤ X, we came to realize that mRNA accessibility is a major issue that should be addressed experimentally as a pre-processing step prior to sequence design. To measure base-pairing accessibility directly, we developed a simple cost-effective method, termed “RNAhyb”, that measures hybridization between a full-length mRNA and individual 20-nt ^32^P-labeled DNA probes that each bind to a different subsequence along the mRNA (see Section S6). We discovered that most 20-nt windows within mRNA Y are relatively inaccessible (<10% hybridization yield) and that the most accessible regions permit hybridization yields of 20-40% (Figure 4). Using RNAhyb, we identified a 150-nt window of mRNA Y that is relatively accessible. We then provided NUPACK with this accessible window as a sequence constraint in place of full-length mRNA Y.

The resulting NUPACK sequence design yielded an scRNA A · B that successfully detected full-length mRNA input Y. However, fixing the mRNA accessibility issue revealed a new degradation issue, as the functional scRNA A · B for reverse logic Y ┤ X was rapidly degraded by RNases in the lysate (Figure S15). Interestingly, we had not observed this degradation issue to nearly the same degree with the forward logic scRNA (Figure S16). To this point, both in previous work^8^ and for the forward logic X ┤ Y, we had used a fully modified 2′OMe-RNA A strand to prevent Dicer processing of scRNA A B, but an unmodified B strand to permit Dicer processing of shRNA output B. Now, for the reverse logic Y ┤ X, we found that B was resistant to degradation as a monomer shRNA but susceptible to degradation as part of dimer scRNA A B. To simultaneously prevent degradation of A B and maintain Dicer cleavage of shRNA B, we tested a variety of chemical modifications to B. Conveniently, we discovered that using a 1-nt 2′OMe-RNA cap at either end of the B strand, in conjunction with 20OMe-RNA for all of strand A yielded the desired properties for the reverse mechanism (Figure S15) as well as for the forward mechanism (Figure S16).

**Figure 4.**
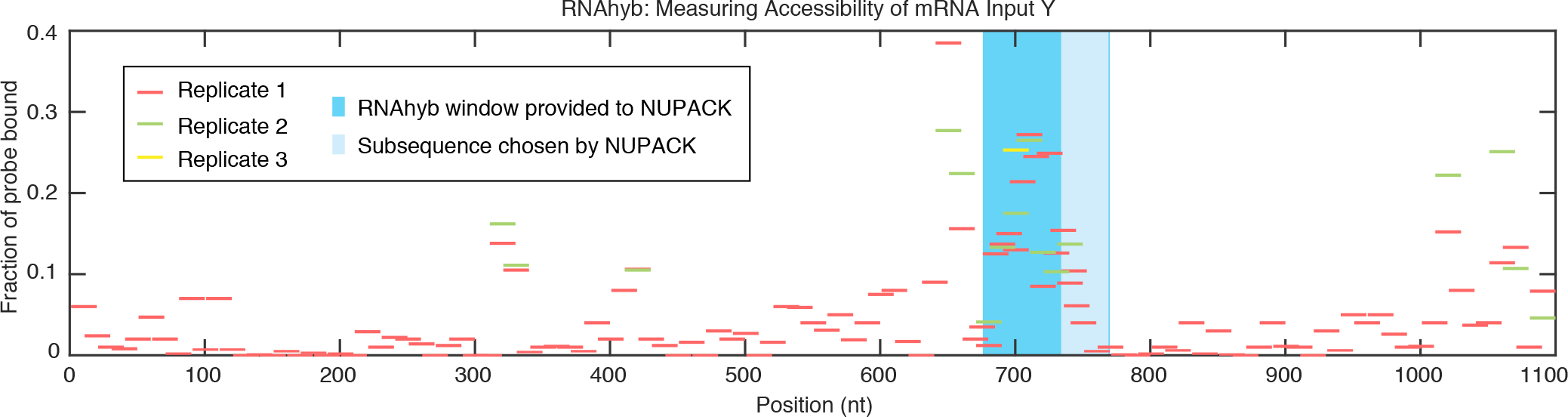
RNAhyb: experimental determination of mRNA accessibility. 20-nt DNA probe at 2.5 nM (labeled with ^32^P), mRNA input Y at 10 nM. Total of 118 probes tested at 10 nt intervals along mRNA input Y to identify an accessible window for engineering reverse logic mechanism (Y ┤ X).

**Figure 5.**
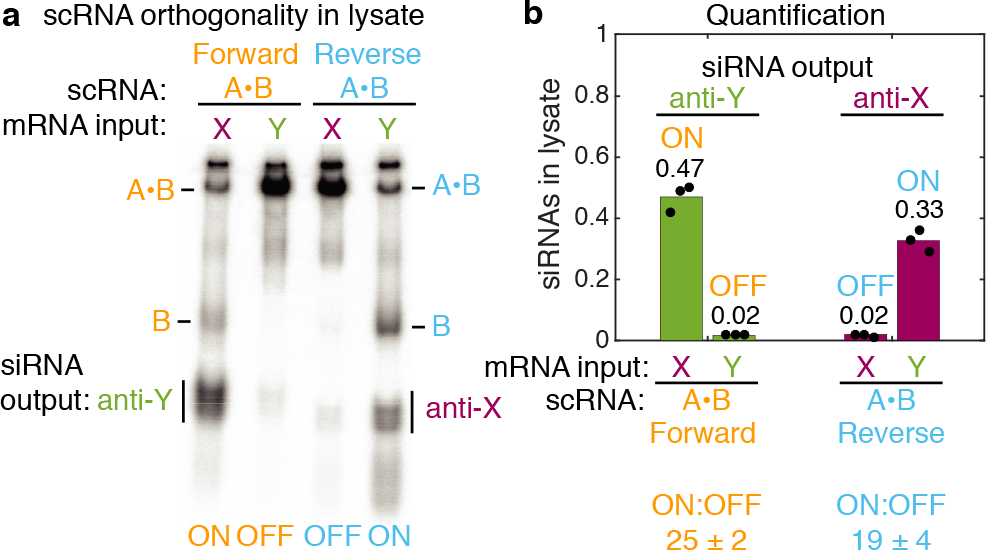
Signal transduction using orthogonal scRNAs in human cell lysate. (a) Conditional siRNA production in the presence of cognate mRNA input (ON state) or non-cognate mRNA input (OFF state) for forward logic (X ┤ Y: cognate mRNA input X, siRNA output anti-Y, non-cognate mRNA input Y) and reverse logic (Y a X: cognate mRNA input Y, siRNA output anti-X, noncognate mRNA input X). Wildtype lysate containing endogenous Dicer. (b) Quantification of siRNA output bands in panel (a). The ON:OFF ratio is ≈20 for both forward and reverse scRNAs. Normalized signal (siRNA signal/total lane signal) for three replicate gels.

Leveraging the mRNA accessibility sequence constraints provided by RNAhyb in combination with the new scRNA chemical modifications yields the reverse mechanism data shown in Figure 3b. This scRNA executes the logic “if Y then not X” (Y ┤ X) with nearly the same characteristics as the initial X ┤ Y mechanism. In the OFF state, scRNA A B produces minimal shRNA B (lane 1) or siRNA output anti-X (lane 2). In the ON state, scRNA A B incubated with full-length mRNA Y produces shRNA B (lane 3), which is cleaved by Dicer into siRNA output anti-X (lane 4). Blocking the nucleation site on mRNA Y restores the OFF state (lanes 5 and 6), demonstrating that scRNA signal transduction is triggered by the intended subsequence of mRNA input Y.

Programmability, the ability to redesign scRNAs to interact with mRNAs X and Y of choice, and orthogonality, the ability to engineer scRNAs that operate independently without crosstalk, are key conceptual goals of dynamic RNA nanotechnology. Both properties are highlighted in a side-by-side
demonstration of the forward system (X ┤ Y) and reverse system (Y ┤ X) in Figure 5a. Each system produces a strong ON state when presented with its cognate mRNA input (X for X ┤ Y; Y for Y ┤ X), and a clean OFF state when presented with the wrong input (Y for X ┤ Y; X for Y ┤ X) even though both scRNAs are constrained by the sequences of both X and Y. Quantification of the siRNA output bands for both forward and reverse systems reveals a strong OFF/ON conditional response with low crosstalk, corresponding to a ≈20-fold increase in production of the siRNA output in response to the cognate vs non-cognate full-length mRNA input.

In the long run, we are working towards the goal of establishing a technology development pipeline for scRNA signal transduction in living organisms. Starting with a functional goal for programmable conditional regulation (e.g., cell-selective gene silencing, gene activation, or cell death), the first step is to decide which programmable input and endogenous pathway to leverage (Figure 1b), followed by invention of an scRNA signal transduction mechanism exploiting design elements from dynamic RNA nanotechnology (Figure 1a), computational sequence design (Figure 2), and then experimental validation in increasingly complex experimental settings: buffer, cell lysate, cell culture, and finally, living organisms. Here, we established human cell lysate as an scRNA testbed intermediate between the chemically pristine environment of buffer in a test tube and the more challenging compartmentalized setting of the living cell. Radioactive labeling of scRNA B allowed use of a biologically relevant scRNA concentration (2.5 nM) in lysate containing cellular proteins and nucleic acids including RNases. The lysate environment enabled optimization of domain dimensions for scRNAs that were functional in buffer but non-functional in lysate as well as optimization of scRNA chemical modifications to inhibit off-pathway degradation while retaining on-pathway Dicer processing. These design changes are likely also necessary (but probably not sufficient) for scRNAs to be functional in living cells, an environment where pathway interrogation is much more difficult.

While lysate offers the key benefit of eliminating the need for scRNA delivery, this simplicity comes with some drawbacks. Any design challenges related to cellular compartmentalization are lost in lysate. Dilution of cellular components in lysate decreases protein and nucleic acid concentrations below those in the cell, limiting our ability to characterize off-target sequence effects, and likely also contributing to our inability to detect cellular mRNA inputs in lysate. Here, we spiked in vitro transcribed mRNA inputs into the lysate.

mRNA accessibility was another important stumbling block, limiting the utility of multiple scRNAs that performed the intended signal transduction when presented with a short RNA input, but failed to detect the same short sequence when it was embedded in a full-length mRNA. To overcome this difficulty, we developed the RNAhyb assay to directly characterize the most accessible windows within an RNA input as a pre-processing step prior to sequence design.

Automated scRNA sequence design was performed using the NUPACK multistate designer subject to biological sequence constraints as well as those imposed by the dynamic scRNA reaction pathway. With this approach, we achieved conditional siRNA production with a ≈20-fold OFF/ON response in human cell lysate for both the forward molecular logical X ┤ Y and the reverse molecular logic Y ┤ X, where X and Y are full-length mRNAs. These results illustrate the programmability and conditionality that make scRNA signal transduction an alluring goal.

If the challenges of engineering and delivering scRNAs in living organisms can be overcome, they offer the potential to serve as powerful new research tools, leveraging diverse endogenous pathways including RNAi^5–9^ and CRISPR/Cas9.^19, 20^ For example, conditional gene silencing (“if gene X is transcribed, silence independent gene Y”) would
probe genetic necessity, conditional gene activation (“if gene X is transcribed, activate independent gene Y”) would probe genetic sufficiency, and conditional cell death (“if gene X is transcribed, induce apoptosis”) would probe developmental compensation. In each case, conditional regulation would be mediated by scRNAs that interact and change conformation to transduce between detection of programmable input X and activation of the desired regulatory output function. By selecting a transcript X with desired spatial and temporal expression profiles, the regulatory function could be restricted to a desired cell type, tissue, or organ within a model organism. To shift conditional regulation to a different tissue or developmental stage, the scRNAs would be reprogrammed to recognize a different input X. The same molecular logic would have attractive therapeutic potential, with X as a programmable disease marker and the downstream regulatory output chosen to be an independent therapeutic pathway, enabling selective treatment of diseased cells. Because of this research and therapeutic potential, dynamic RNA nanotechnology for scRNA signal transduction merits significant engineering effort from the molecular programming and synthetic biology research communities.

## METHODS SUMMARY

Sequence design was performed using the reaction pathway designer within NUPACK.^13, 14^ Oligonucleotides (RNA and 20OMe-RNA) were synthesized and RNase-free HPLC purified by IDT. Target mRNAs were in vitro transcribed. Cell lysates were made by sonication of HEK 293T cells. Dicer-depleted lysates were generated by knocking down Dicer using an siRNA pool prior to lysis and Dicer immunodepletion following lysis. ^32^P was incorporated in the backbone of B to enable discrimination of scRNA reactants, intermediates, and products from lysate nucleic acids, as well as to enable gel assays at low concentration. scRNA A · B was PAGE purified as a duplex. scRNA signal transduction was tested by incubating 2.5 nM scRNAs and 10 nM short RNA input or full-length mRNA input in 1× Buffer D or 1.25 *μ*g/*μ*L lysate at 37 °C for 4 h. For experiments with a blocked nucleation site, the mRNA input was pre-incubated with blocker L. Reactions were separated by native PAGE and imaged via autoradiography. RNAhyb was used to assay mRNA accessibility using 20-nt ^32^P-labeled DNA probes at 10-nt intervals along the mRNA input.

## ASSOCIATED CONTENT

Detailed methods, RNAhyb protocol, sequences, motivating failures, additional signal transduction studies for forward and reverse mechanisms, RNAhyb studies, chemical modification studies.

## AUTHOR INFORMATION

### Notes

The authors declare the following competing financial interests: US patents.

## ACKNOWLEDGMENTS

We thank K. Sakurai and J.J. Rossi for advice on preparation of human cell lysate with functional Dicer, B.R. Wolfe and N.J. Porubsky for assistance with reaction pathway engineering using NUPACK, and P.D. Carlson and J.B. Lucks for discussions on profiling mRNA accessibility. This work was funded by the National Institutes of Health (5R01CA140759), by the National Science Foundation Molecular Programming Project (NSF-CCF-1317694), by the Gordon and Betty Moore Foundation (GBMF2809), by the Rosen Bioengineering Center at Caltech, by a Professorial Fellowship at Balliol College (University of Oxford), and by the Eastman Visiting Professorship at the University of Oxford.

